# MVP: predicting pathogenicity of missense variants by deep learning

**DOI:** 10.1101/259390

**Authors:** Hongjian Qi, Chen Chen, Haicang Zhang, John J. Long, Wendy K. Chung, Yongtao Guan, Yufeng Shen

## Abstract

Accurate pathogenicity prediction of missense variants is critical to improve power in genetic studies and accurate interpretation in clinical genetic testing. Here we describe a new prediction method, MVP, which uses a deep learning approach to leverage large training data sets and many correlated predictors. Using cancer mutation hotspots and *de novo* germline mutations from developmental disorders for benchmarking, MVP achieved better performance in prioritizing pathogenic missense variants than previous methods.

## Main Text

Missense variants are the most common type of coding genetic variants and are a major class of genetic risk across a broad range of common and rare diseases. Previous studies have estimated that there is a substantial contribution from *de novo* missense mutations to structural birth defects^1–3^ and neurodevelopmental disorders^4–6^. However, only a small fraction of missense *de novo* mutations are pathogenic^4^. As a result, the statistical power of detecting individual risk genes based on missense variants or mutations is limited^7^. In clinical genetic testing, many of missense variants in well-established risk genes are classified as variants of uncertain significance, unless they are highly recurrent in patients. Previously published *in silico* prediction methods have facilitated the interpretation of missense variants, such as CADD^8^, VEST3^9^, metaSVM^10^, M-CAP^11^, and REVEL^12^. However, based on recent *de novo* mutation data, they all have limited performance with low positive predictive value (Supplementary Table S1), especially in non-constrained genes (defined as ExAC^13^ pLI<0.5).

Here we hypothesize that missense variant pathogenicity prediction can be improved in a few dimensions. First, conventional machine learning approaches have limited capacity to leverage large amount of training data compared to recently developed deep learning methods^14^. Second, databases of pathogenic variants curated from the literature are known to have a substantial frequency of false positives^15^, which are likely caused by common issues across databases and therefore introduce inflation of benchmark performance. Developing new benchmark data and methods can help to assess and improve real performance. Finally, previous methods do not consider gene dosage sensitivity^13, 16^, which can modulate the pathogenicity of deleterious missense variants, as hypomorphic variants are pathogenic only in dosage sensitive genes^6^. With recently published metrics of mutation intolerance, it is now feasible to consider gene dosage sensitivity in predicting pathogenicity. Based on these ideas, we developed a new method, MVP, to improve missense variant pathogenicity prediction.

MVP uses many correlated predictors, which can be broadly grouped into two categories (Supplementary Table S2): (a) “raw” features computed at different scales, per base pair (e.g. amino acid constraint score and conservation), per local context (e.g. protein structure and modification) as well as per gene (e.g. gene mutation intolerance, sub-genic regional depletion of missense variants ^17^); (b) deleteriousness scores from selected previous methods. We reason that the variants in constrained genes (ExAC pLI≥0.5) and non-constrained genes may have different modes of action of pathogenicity, therefore, trained our models for the two gene sets separately. We included 38 features for the constrained gene model, and 21 features for the non-constrained gene where we removed most published prediction methods features due to limited prediction accuracy (Supplementary Table S1, S2).

MVP uses a deep residual neural network (ResNet)^18^ model. There are two layers of residual blocks, consisting of convolutional filters and activation layers, and two fully connected layers with sigmoid output (Supplementary Fig. S1). The convolutional filters can exploit spatial locality by enforcing a local connectivity pattern between “neurons” of adjacent layers and identify nonlinear interactions at higher levels of the network. To take advantage of this characteristic, we ordered the predictors based on their correlation, as highly correlated predictors are clustered together (Supplementary Fig. S2). Notably, some protein-related predictors are weakly correlated with previous scores, suggesting that they may include additional information and can help improve the overall prediction accuracy. For each missense variant, we defined MVP score by the rank percentile of the ResNet’s raw sigmoid output relative to all 76 million possible missense variants.

We obtained large curated datasets of pathogenic variants as positives and random rare missense variants from population data as negatives for training (Supplementary Table S3). Using 6-fold cross-validation on the training set (Supplementary Fig. S3), MVP achieved mean area under the curve (AUC) of 0.99 in constrained genes and 0.97 in non-constrained genes.

To evaluate predictive performance of the MVP and compare it with other methods, we evaluated the performance in an independent curated testing dataset from VariBench^10, 19^ (Supplementary Fig. S4). MVP outperformed all other scores with an AUC of 0.96 and 0.92 in constrained and non-constrained genes, respectively. A few recently published methods (REVEL, M-CAP, VEST3, and metaSVM) were among the second-best predictors and achieved AUC around 0.9.

Systematic false positives caused by similar factors across training and VariBench data sets could inflate the performance in testing. To address this issue, we obtained two additional types of data for further evaluation. First, we compiled cancer somatic mutation data, including missense mutations located in inferred hotspots based on statistical evidence from a recent study^20^ as positives, and randomly selected variants from DiscovEHR^21^ database as negatives. In this dataset, the performance of all methods decreased, but MVP still achieved the best performance of AUC of 0.91 and 0.85 in constrained and non-constrained genes, respectively (Fig. 1). We observed that methods using HGMD or UniProt in training generally have greater performance drop than others (Supplementary Table S4, Fig. S5, Supplementary notes).

**Figure. 1:**
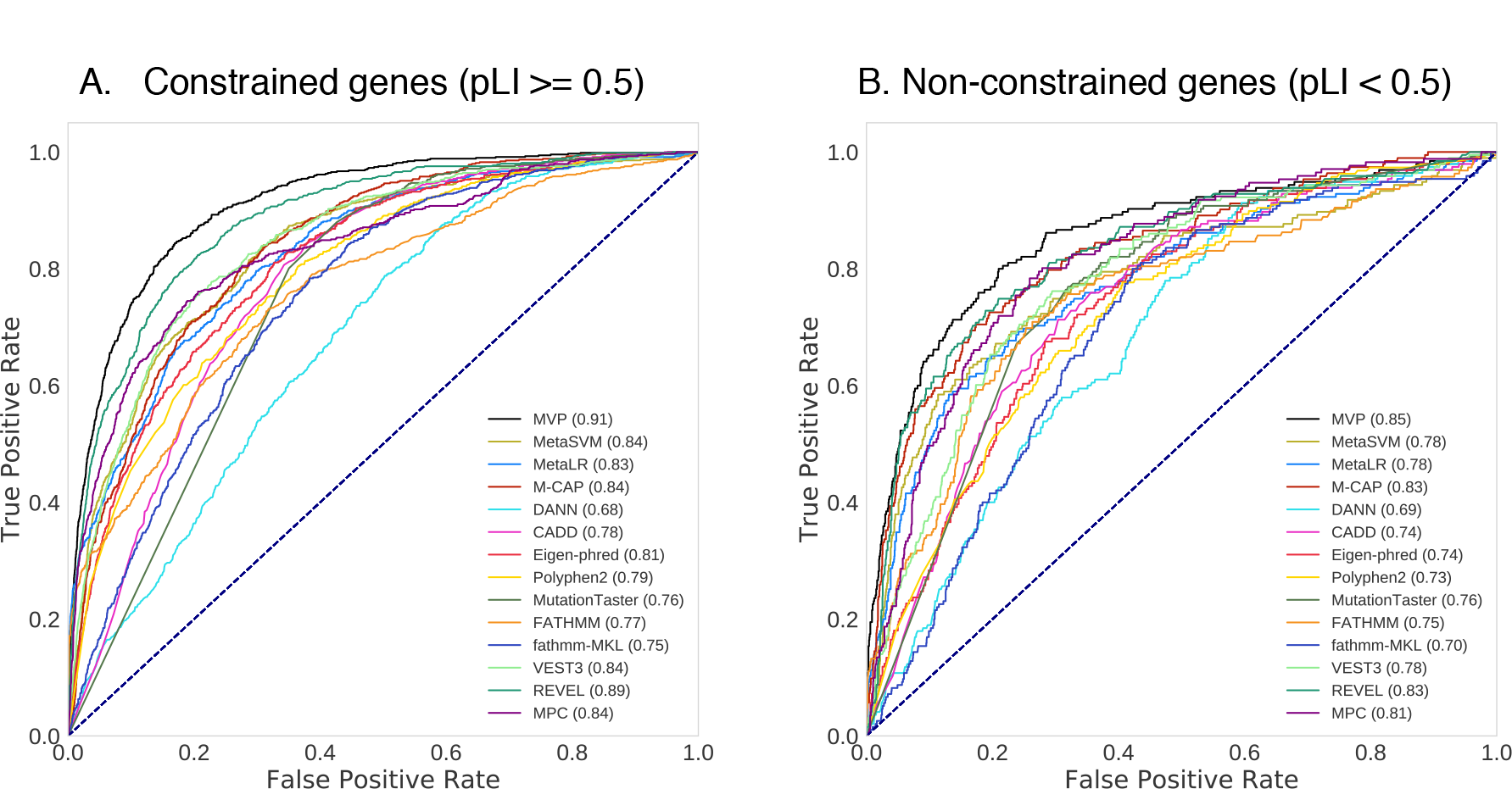
ROC curves for existing prediction scores and MVP scores of cancer somatic mutation data sets. (A) Constrained genes: evaluation of 699 cancer mutations located in hotspots from 150 genes, and 6989 randomly selected mutations from DiscovEHR database excluding mutations used in training. (B) Non-constrained genes: evaluation of 177 cancer mutations located in hotspots from 55 genes and 1782 randomly selected mutations from DiscovEHR database excluding mutations used in training. The performance of each method is evaluated by the ROC curve and AUC score indicated in parenthesis. Higher AUC score indicates better performance.

To investigate the contribution of features to MVP predictions, we performed cross-one-group-out experiments and used the differences in AUC as an estimation of feature contribution (Supplementary Fig. S6). We found that in constrained gene, conservation scores and published deleteriousness predictors have relatively large contribution, whereas in non-constrained genes, protein structure and modification features and published predictors are most important.

Second, to test the utility in real genetic studies, we obtained germline *de novo* missense variants (DNMs) from 2645 cases in a congenital heart disease (CHD) study^2^, 3953 cases in autism spectrum disorder (ASD) studies^2, 4, 5^, and DNMs from 1911 controls (unaffected siblings) in Simons Simplex Collection^2, 4, 5^. Since genes with cancer mutation hotspots are relatively well studied in both constrained and non-constrained gene sets, assessment using *de novo* mutations can provide additional insight with less bias (Supplementary Table S5). Because the true pathogenicity of most of the *de novo* mutations is unknown, we cannot directly evaluate the performance of prediction methods. To address this issue, we calculated the enrichment rate of predicted pathogenic DNMs by a method with a certain threshold in the cases compared to the controls, and then estimated precision and the number of true risk variants (Methods), which is a proxy of recall since the total number of true positives in all cases is a (unknown) constant independent of methods. We compared the performance of MVP to other methods by estimated precision and recall-proxy (Fig. 2). Based on the optimal thresholds of MVP in cancer hotspot ROC curves, we used a score of 0.7 in constrained genes and 0.75 in non-constrained genes to define pathogenic DNMs (Fig. S7). In constrained genes, we observed an enrichment of 2.2 in CHD and an enrichment of 1.9 in ASD (Supplementary Table S6, S7), achieving estimated precision of 0.55 and 0.47 (Fig. 2A and 2D), respectively. This indicates that about 50% of the MVP-predicted pathogenic DNMs contribute to the diseases. In non-constrained genes, we observed an enrichment of 1.9 in CHD and 1.4 in ASD (Supplementary Table S6, S7), respectively, and 0.32 and 0.28 in estimated precision (Fig. 2B and 2E). In all genes combined, MVP achieved an estimated precision of 40% for both CHD and ASD (Fig. 2C and 2F). The next best methods reached 25% (M-CAP) and 20% (MPC^17^ and REVEL) given the same recall-proxy for CHD and ASD, respectively (Supplementary Table S6, S7). Furthermore, the estimated precision of MVP with DNMs at optimal threshold is much closer to the expected precision based on ROC of cancer hotspots data than the value from VariBench data (Supplementary Figure S8 and Notes), supporting that there is less performance inflation in testing using cancer data.

**Figure. 2:**
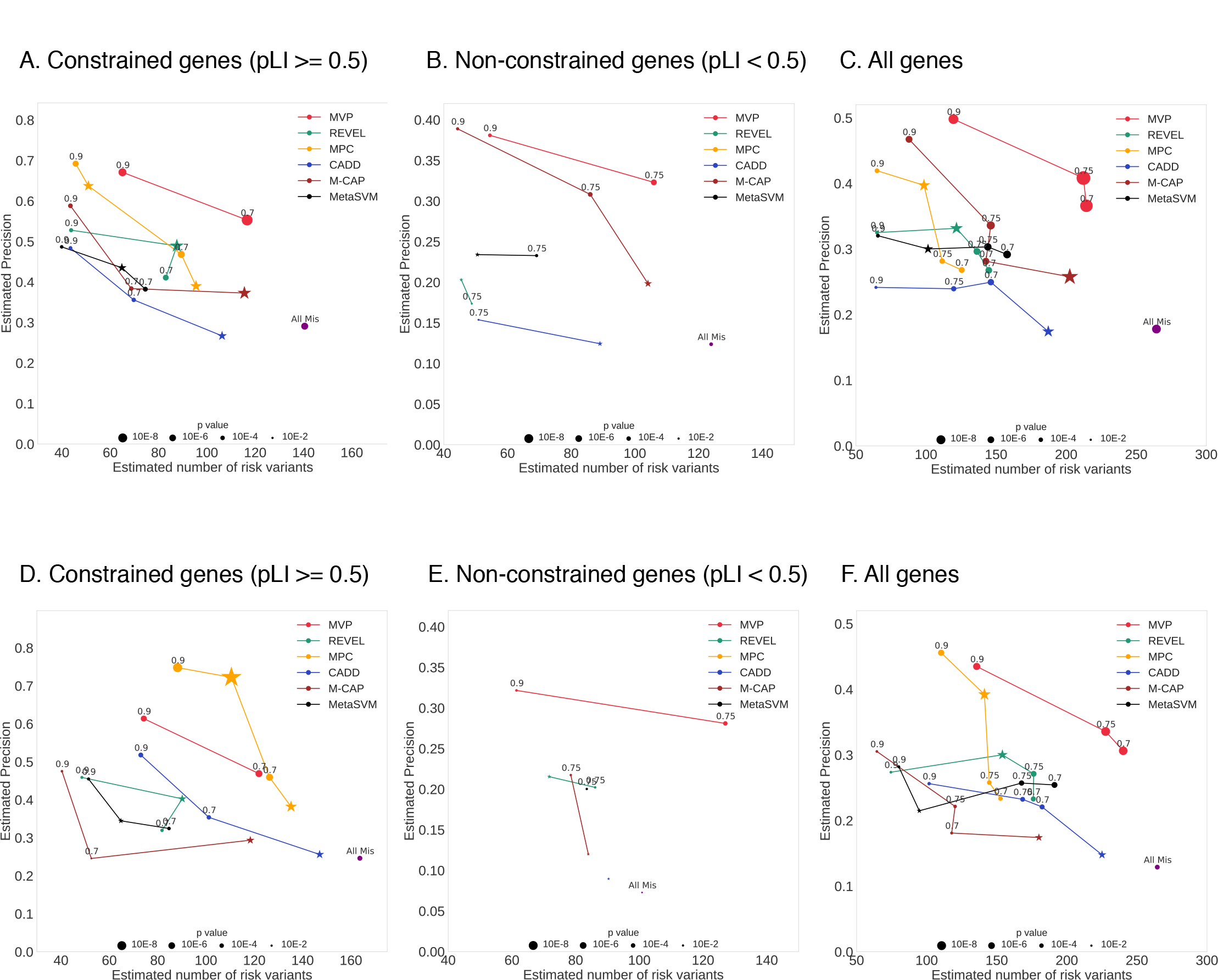
Comparison of MVP and previously published methods using *de novo* missense mutations from CHD and ASD studies by precision-recall-proxy curves. Numbers on each point inchcate rank percentile thresholds, star points indicate thresholds recommended by publications. The positions of “All Mis” points are estimated from all missense variants in the gene set without using any pathogenicity prediction method. The point size is proportional to -log (p-value). P-value is calculated by binomial test, only points with p value less than 0.05 are shown. (A, B, C) Performance in CHD DNMs in constrained genes, non-constrained genes, and all genes, respectively. (D, E, F) Performance in ASD DNMs in constrained genes, non-constrained genes, and all genes, respectively.

Previous studies have estimated that deleterious *de novo* coding mutations, including loss of function variants and damaging missense variants, have a small contribution to isolated CHD^2^. Here, we used MVP to revisit this question. With the definition of damaging DNMs in Jin et al 2017^2^ (based on metaSVM^10^), the estimated contribution of deleterious *de novo* coding mutations to isolated CHD is about 4.3%. With MVP score of 0.75, the estimation is 7.8%(95% CI = [6.5%, 9.1%]), nearly doubling the previous estimate (Supplementary Table S8, S9).

In summary, we developed a new method, MVP, to predict pathogenicity of missense variants. MVP is based on residual neural networks, a supervised deep learning approach, and was trained using a large number of curated pathogenic variants from clinical databases, separately on constrained genes and non-constrained genes. Using cancer mutation hotspots and *de novo* mutations from CHD and ASD, we showed that MVP achieved overall better performance than published methods, especially in non-constrained genes. Nevertheless, the fraction of pathogenic variants among *de novo* missense variants in non-constrained genes is low in both CHD and ASD, leading to relatively poor performance by all methods. MVP achieved substantially better performance than other methods in these genes, partly attributed to inclusion of protein structure-based predictors (Supplementary Figure S6B). Further improvement in protein structure prediction and the utilization of protein structure in the model ^22^ would be the key to improve MVP. Finally, all methods are limited by the size and the potentially high false positive rate of the training data. Systematic efforts such as ClinVar^23^ will eventually produce better training data to improve prediction performance.

## ULRs

Software and data for implementing MVP are available from https://github.com/ShenLab/missense;

Precomputed MVP pathogenicity score for all possible missense variants in canonical transcripts on human hg19 can be downloaded from: https://www.dropbox.com/s/bueatvqnkvqcb54/MVPscoreshg19.txt.bz2?dl=0

## Methods and materials

### Training data sets

We compiled 22,390 missense mutations from Human Gene Mutation Database Pro version 2013 (HGMD)^24^ database under the disease mutation (DM) category, 12,875 deleterious variants from UniProt^10, 25^, and 4,424 pathogenic variants from ClinVar database^23^ as true positive (TP). In total, there are 32,074 unique positive training variants. The negative training sets include 5,190 neutral variants from Uniprot^10, 25^, randomly selected 42,415 rare variants from DiscovEHR database^21^, and 39,593 observed human-derived variants^8^. In total, there are 86,620 unique negative training variants (Supplementary Table S3).

### Testing data sets

We have three categories of testing data sets (Supplementary Table S3). The three categories are: (a) Benchmark data sets from VariBench ^10, 19^ as positives and randomly selected rare variants from DiscovEHR database^21^ as negatives; (b) cancer somatic missense mutations located in hotspots from recent study^26^ as positives and randomly selected rare variants from DiscovEHR database^21^ as negatives; (c) and *de novo* missense mutation data sets from recent published exome-sequencing studies^2, 4, 5^. All variants in (a) and (b) that overlap with training data sets were excluded from testing.

We tested the performance in constrained genes (ExAC pLI ≥ 0.5) and non-constrained gene (ExAC pLI < 0.5)^13^ separately.

To focus on rare variants with large effect, we selected ultra-rare variants with MAF <10^−4^ based on gnomAD database to filter variants in both training and testing data sets. We applied additional filter of MAF < 10^−6^ for variants in constrained genes in both cases and controls for comparison based on a recent study ^17, 27^.

## Features used in MVP model

MVP uses many correlated features as predictors (Supplementary Table S2). There are six categories: (1) local context: GC content within 10 flanking bases on the reference genome; (2) amino acid constraint, including blosum62^28^ and pam250^29^; (3) conservation scores, including phyloP 20way mammalian and 100way vertebrate^30^, GERP++^31^, SiPhy 29way^32^, and phastCons 20way mammalian and 100way vertebrate^33^; (4) Protein structure, interaction, and modifications, including predicted secondary structures^34^, number of protein interactions from the BioPlex 2.0 Network^35^, whether the protein is involved in complexes formation from CORUM database^36^, number of high-confidence interacting proteins by PrePPI ^37^, probability of a residue being located the interaction interface by PrePPI (based on PPISP, PINUP, PredU), predicted accessible surface areas were obtained from dbPTM^38^, SUMO scores in 7-amino acids neighborhood by GPS-SUMO ^39^, phosphorylation sites predictions within 7 amino acids neighborhood by GPS3.0^40^, and ubiquitination scores within 14-amino acids neighborhood by UbiProber ^41^; (5) Gene mutation intolerance, including ExAC metrics^13^ (pLI, pRec, lof_z) designed to measure gene dosage sensitivity or haploinsufficiency, RVIS^42^, probability of causing diseases under a dominant model “domino”^43^, average selection coefficient of loss of function variants in a gene “s_het” ^44^, and sub-genic regional depletion of missense variants ^17^; (6) Selected deleterious or pathogenicity scores by previous published methods obtained through dbNSFPv3.3a^45^, including Eigen^46^, VEST3^9^, MutationTaster^47^, PolyPhen2 ^48^, SIFT ^49^, PROVEAN^50^, fathmm-MKL ^51^, FATHMM ^51^, MutationAssessor^52^, and LRT^53^.

For consistency, we used canonical transcripts to define all possible missense variants^17^. Missing values of protein complex scores are filled with 0 and other features are filled with −1.

Since pathogenic variants in constrained genes and non-constrained genes may have different mode of action, we trained our models on constrained and non-constrained variants separately with different sets of features (38 features used in constrained model, 21 features used in non-constrained model, Supplementary Table S2).

## Deep learning model

MVP is based on a deep residual neural network model (ResNet)^18^ for predicting pathogenicity using the predictors described above. To preserve the structured features in training data, we ordered the features according to their correlations (Supplementary Fig. S2). The model (Supplementary Figure S1) takes a vector of the ordered features as input, followed by a convolutional layer of 32 kernels with size 3 x 1 and stride of 1, then followed by 2 computational residual units, each consisting of 2 convolutional layers of 32 kernels with size 3 x 1 and a ReLU^54^ activation layer in between. The output layer and input layer of the residual unit is summed and passed on to a ReLU activation layer. After the two convolutional layers with residual connections, 2 fully connected layers of 320 x 512 and 512 x1 are used followed by a sigmoid function to generate the final output^55^.

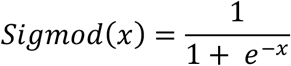

(Supplementary Fig. S1).

In training, we randomly partitioned the synthetic training data sets into two parts, 80% of the total training sets for training and 20% for validation. We trained the model with batch size of 64, used adam^56^ optimizer to perform stochastic gradient descent^57^ with logarithmic loss between the predicted value and true value. After one full training cycle on the training set, we applied the latest model weights on validation data to compute validation loss.

To avoid over fitting, we used early stopping regularization during training. We computed the loss in training data and validation data after each training cycle and stopped the process when validation loss is comparable to training loss and do not decrease after 5 more training cycle, and then we set the model weights using the last set with the lowest validation loss. We applied the same model weights on testing data to obtain MVP scores for further analysis.

## Previously published methods for comparison

We compared MVP score to 13 previously published prediction scores, namely, M-CAP^11^, DANN^58^, Eigen^46^, Polyphen2^48^, SIFT^49^, MutationTaster^47^, FATHMM^51^, REVEL^12^, CADD^8^, metaSVM^10^, metaLR^10^, VEST3^9^, and MPC^17^.

## Normalization of scores using rank percentile

For each method, we first obtained predicted scores of all possible rare missense variants in canonical transcripts, and then sort the scores and converted the scores into rank percentile. Higher rank percentile indicates more damaging, e.g., a rank score of 0.75 indicates the missense variant is more likely to be pathogenic than 75% of all possible missense variants.

## ROC curves

We plotted Receiver operating characteristic (ROC) curves and calculated Area Under the Curve (AUC) values in training data with 6-fold cross validation (Supplementary Fig. S3), and compared MVP performance with other prediction scores in curated benchmark testing datasets (Supplementary Fig. S4) and cancer hotspot mutation dataset (Fig. 2). For each prediction method, we varied the threshold for calling pathogenic mutations in a certain range and computed the corresponding sensitivity and specificity based on true positive, false positive, false negative and true negative predictions. ROC curve was then generated by plotting sensitivity against 1 - specificity at each threshold.

## Optimal points based on ROC curves

We define the optimal threshold for MVP score as the threshold where the corresponding point in ROC curve has the largest distance to the diagonal line (Supplementary Figure S7). Based on the true positive rate and false positive rate at the optimal points in ROC curves, we can estimate the precision and recall in *de novo* precision-recall-proxy curves (Supplementary Figure S8 and Supplementary Notes).

## Precision-recall-proxy curves

Since *de novo* mutation data do not have ground truth, we used the excess of predicted pathogenic missense *de novo* variants in cases compared to controls to estimate precision and proxy of recall. For various thresholds of different scores, we can calculate the estimated number of risk variants and estimated precision based on enrichment of predicted damaging variants in cases compared to controls. We adjusted the number of missense *de novo* mutation in controls by the synonymous rate ratio in cases verses controls, assuming the average number of synonymous as the data sets were sequenced and processed separately) (Table S10), which partly reduced the signal but ensures that our results were not inflated by the technical difference in data processing.

Denote the number of cases and controls as *N*_*1*_ and *N*^*0*^, respectively; the number of predicted pathogenic *de novo* missense variants as *M*_*1*_ and *M*^*0*^, in cases and controls, respectively; the rate of synonymous *de novo* variants as *S*_*1*_ and *S*_*0*_, in cases and controls, respectively; technical adjustment rate as *α*; and the enrichment rate of variants in cases compared to controls as *R*.

We first estimate α by:

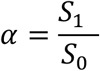

Then assuming the rate of synonymous de novo variants in cases and controls should be identical if there is no technical batch effect, we use α to adjust estimated enrichment of pathogenic de novo variants in cases compared to the controls by:

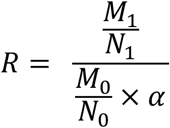

Then we can estimate number of true pathogenic variants 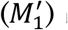 by:

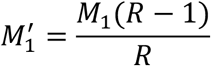

And then precision by:

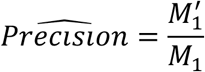

## Acknowledgements

We thank Na Zhu, Ben Lai, Itsik Pe’er, Emily Gao, and Jiayao Wang for helpful discussions. We thank Pediatric Cardiac Genomics Consortium (PCGC) investigators for CHD data access, and the patients and their families for their generous contribution to the PCGC study and to the Simons Simplex Collection study families and principal investigators (A.L. Beaudet, R. Bernier, J. Constantino, E.H. Cook, Jr, E. Fombonne, D. Geschwind, D.E. Grice, A. Klin, D.H. Led-better, C. Lord, C.L. Martin, D.M. Martin, R. Maxim, J. Miles, O. Ousley, B. Peterson, J. Piggot, C. Saulnier, M.W. State, W. Stone, J.S. Sutcliffe, C.A. Walsh, and E. Wijsman) and the coordinators and staff at the SSC clinical sites.

